# Muramyl endopeptidase Spr contributes to intrinsic vancomycin resistance in *Salmonella enterica* serovar Typhimurium

**DOI:** 10.1101/274886

**Authors:** Kim Vestö, Douglas L Huseby, Snygg Iina, Helen Wang, Diarmaid Hughes, Mikael Rhen

## Abstract

The impermeability barrier provided by the outer membrane of enteric bacteria, a feature lacking in Gram-positive bacteria, plays a major role in maintaining resistance to numerous antimicrobial compounds and antibiotics. Here we demonstrate that mutational inactivation of *spr*, coding for a muramyl endopeptidase, significantly sensitizes *Salmonella enterica* serovar Typhimurium to vancomycin without any accompanying apparent growth defect or outer membrane destabilization. A similar phenotype was not achieved by deleting the *mepA, pbpG, nlpC, yebA* or *yhdO* genes coding for functional homologues to Spr. The *spr* mutant revealed increased autolytic behavior in response to not only vancomycin, but also to penicillin G, an antibiotic for which the mutant displayed a wild-type MIC. A screen for suppressor mutations of the *spr* mutant phenotype revealed that deletion of *tsp* (prc), encoding a periplasmic carboxypeptidase involved in processing Spr and PBP3, restored intrinsic resistance to vancomycin and reversed the autolytic phenotype of an *spr* mutant. Our data suggest that Spr contributes to intrinsic antibiotic resistance in S. Typhimurium without directly affecting the outer membrane permeability barrier. Furthermore, our data suggests that compounds that target specific cell wall endopeptidases might have the potential to expand the activity spectrum of traditional Gram-positive antibiotics.

## INTRODUCTION

Peptidoglycan (murein) constitutes a main component of the bacterial cell wall. It is composed of repeated N-actetylglucosamine (GlcNAc) and N-acetylmuramic acid (MurNAc) disaccharide units, cross-linked by peptide bridges. The synthesis of this mesh in turn is the target of several classes of antibiotics, such as the β-lactams and glycopeptides. The outer membrane of Gram-negative enteric bacteria, due to its relative impermeability, provides an intrinsic resistance barrier against many large molecular compounds, including selected antibiotics such erythromycin (Delcour 2009; Nikaido and Vaara 1985), novobiocin (Anderle et al. 2008), rifampicin and the glycopeptide vancomycin (Krishnamoorthy et al. 2013; Weeks et al. 2010).

Peptidoglycan maintains bacterial shape, septum formation at the point of cell division, and cell integrity upon internal turgor stress. To accomplish this, peptidoglycan not only needs to assemble, but also disassemble, in order to facilitate changes in size and outline of the cell wall in growing bacteria. In this, several periplasmic amidases, endopeptidases and glycosylases participate together with penicillin-binding proteins (PBPs), themselves transpeptidases, to shape the peptidoglycan (Sauvage et al. 2008). While the PBPs have the major responsibility of catalyzing the formation of interpeptide bridges between overlapping GlcNAc-MurNAc polymers, the murein endopeptidases are tasked with cleaving interpeptide bridges to facilitate incorporation of new GlcNAC-MurNAc polymers into the growing peptidoglycan mesh. The importance of correctly balancing these opposing activities is illustrated by the fact that blocking PBP activity with β-lactams results in autolysis in *Escherichia coli* (Prestidge and Pardee 1957).

The increasing frequency of clinical bacterial isolates producing extended-spectrum β-lactamases is limiting the effectiveness of antibiotics that target cell wall synthesis amongst Gram-negative species (Coque, Baquero, and Canton 2008). The search for new antibiotics to treat Gram-negative bacterial infections would be advanced by a better understanding of bacterial cell wall homeostasis at the level of peptidoglycan. Because it is a genetically amenable bacterium, *E. coli* has been the main focus for studies on the activities of cell wall-modulating enzymes. From these studies a consensus has emerged that, apart from PBP3 (Botta and Park 1981), each of the glycolytic, endopeptic hydrolases and PBPs are individually dispensable for bacterial viability. Accordingly, inactivation of any one (or sometimes more than one) of the genes encoding these enzymes (PBP4: (Denome et al. 1999; Michio Matsuhashi et al. 1977; Meberg et al. 2004), PBP5: (Denome et al. 1999; M Matsuhashi et al. 1979; Nishimura et al. 1980; Spratt 1980), PBP6: (Denome et al. 1999; Broome-Smith and Spratt 1982), PBP6b: (Baquero et al. 1996), PBP7/PBP8: (Denome et al. 1999; Henderson, Templin, and Young 1995), reviewed in: (van Heijenoort 2011)) does not prevent bacterial growth under laboratory conditions. While this might imply a high degree of functional redundancy, it does not exclude the possibility that some or all of these enzymes may have unique functions under other more specific conditions.

A recent study (Singh et al. 2012) confirmed muramyl endopeptidase activity for three additional *E. coli* proteins; Spr, YebA and YdhO, renamed in *E. coli* to MepS, MepM and MepH (Singh et al. 2015). More specifically the study presented data implying that Spr or YebA might represent endopeptidases with less redundant functions in that it was feasible to construct an Δ*spr*Δ*yebA* mutant only in an *E. coli* strain genetically complemented with *spr* (*mepS*) (Singh et al. 2012).

*Salmonella enterica* serovar Typhimurium (*S*. Typhimurium) is a Gram-negative enterobacterium with an increasing antibiotic resistance development in its genus (Klemm et al. 2018; Hong et al. 2016; Angelo et al. 2016). As the S. Typhimurium genome includes genes with high sequence similarity to the *mepS*, *mepM* and *mepH* genes of *E. coli*, and given the potential importance of *mepS* and *mepM* for viability of *E. coli* (Singh et al. 2012), we studied the phenotypes of *S*. Typhimurium mutants in which these genes were deleted, either singly or in combination. We characterized Δ*spr*, Δ*yebA* and Δ*ydhO* mutants in terms of their growth and susceptibility profiles against antimicrobials, and in addition Δ*spr* for autolytic behavior. Our findings highlight Spr as a possible new target for antibacterial treatment in order to sensitize *Salmonella* against Gram-positive-specific antibiotics.

## RESULTS

### Construction of Δ*spr*, Δ*yebA* and Δ*ydhO* mutants in *S*. Typhimurium

To assess any functional similarity of the S. Typhimurium homologues to the *E. coli* MepS, MepM and MepH proteins regarding growth phenotypes we constructed single and all combinations of double deletion mutants of Δ*spr*, Δ*yebA* and Δ*ydhO* in *S*. Typhimurium SR-11, using allelic replacement (for details, see Materials and Methods). All deletions were verified by PCR, where after the allelic replacement the antibiotic resistance cassette was removed. In agreement with observations from *E. coli* (Singh et al. 2012) we were not successful in creating a Δ*spr*Δ*yebA*Δ*ydhO* triple mutant.

### Growth characteristics of mutants

In agreement with observations made in *E. coli* (Singh *et al*. 2012) none of the single deletion mutants revealed any significant difference in the overall shape of their growth curves (Fig. 1A and 1B). All single mutants had a similar logarithmic growth rate, and reached a similar optical density in stationary phase in both LB and TY medium. Even the Δ*yebA*Δ*ydhO* mutant grew like the wild type parental strain. On the other hand, the Δ*spr*Δ*yebA* mutant showed a somewhat decreased rate of replication in TY medium at later stages of the growth slope (Fig. 1B). Still, taken together, these findings suggest a high degree of redundancy for the Spr, YebA and YhdO endopeptidases in *S*. Typhimurium regarding growth in broth cultures.

**Figure 1.**
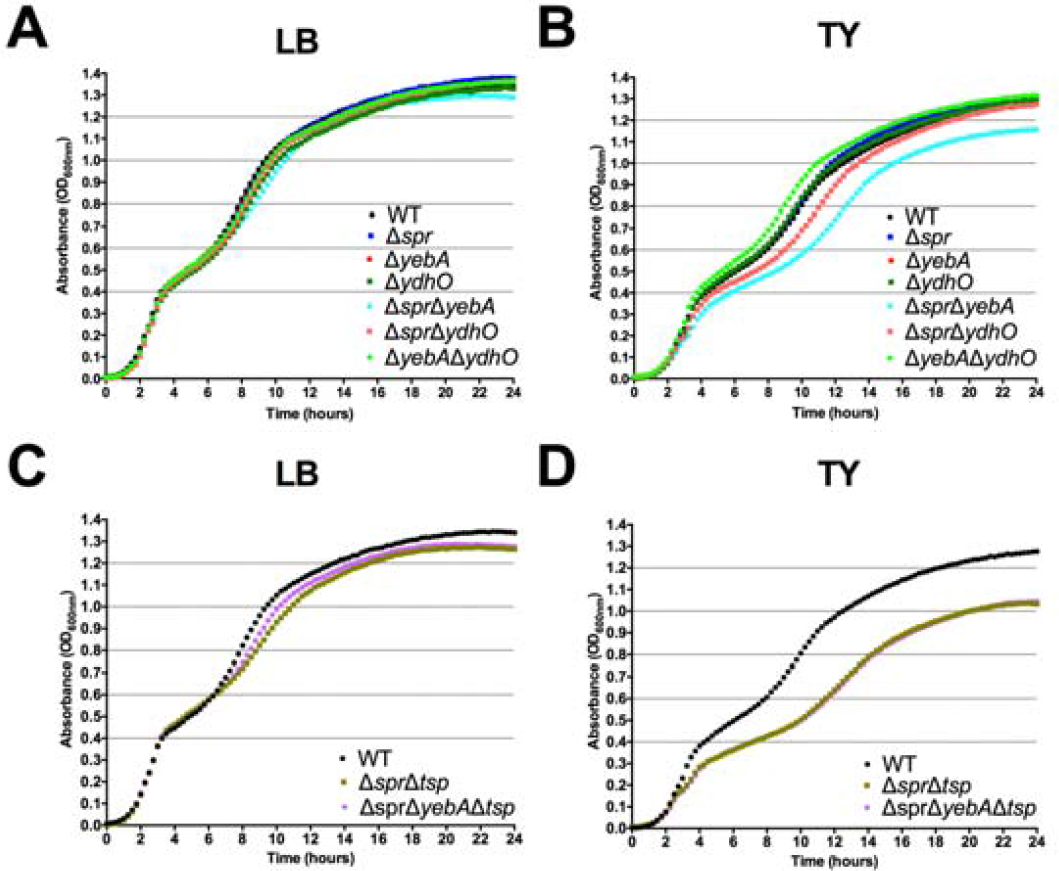
Growth characteristics of S. Typhimurium strains. Bacterial strains were diluted to an OD_600nm_ of 0.01 from overnight LB broth cultures into LB (**A & C**) or TY (**B & D**) broth where after increase in bacterial density was followed for 24 hours.

### Antibiotic and detergent sensitivities

As peptidoglycan turnover and outer membrane synthesis in *E. coli* appear connected (Gray et al. 2015), and as *E. coli* mutants simultaneously lacking several murein hydrolases show evidence of an outer membrane permeability barrier defect (Heidrich et al. 2002), we assessed whether any of the murein endopeptidases Spr, YebA or YhdO were needed for maintaining the outer membrane permeability barrier in *S*. Typhimurium. To achieve this, we screened the panel of our *S*. Typhimurium endopeptidase mutants for possible sensitization to six different antibiotics using the disc diffusion method, as well as for detergent tolerance. The antibiotics tested were penicillin G, polymyxin B, tetracycline, rifampicin, novobiocin, and vancomycin, the latter three for which wild-type *S*. Typhimurium is intrinsically resistant to due to the outer membrane permeability barrier (Sukupolvi et al. 1984). As comparator strains we used wild-type *S*. Typhimurium SR-11, and three isogenic LPS mutants expected to have an outer membrane permeability defect (Sukupolvi *et al*. 1984). Sensitivity testing was conducted using low osmolarity TY medium. The rationale for this was that high salt concentrations reduce the sensitivity of *E. coli* to selected antibiotics (Beggs and Andrews 1976), and that low osmolarity medium would favor expression of the more permeable outer membrane porin OmpF (Jaffe, Chabbert, and Semonin 1982; Harder, Nikaido, and Matsuhashi 1981), thus increasing the probability of detecting any sensitization.

As compared to the wild type the three LPS mutants, Δ*rfaC* (*waaC*), Δ*rfaG* (*waaG*), and Δ*rfaP* (*waaP*) were each sensitized to polymyxin B, novobiocin, rifampicin and vancomycin, but not to tetracycline (Table 1). The Δ*spr* mutant was strongly sensitized to vancomycin (inhibition zone increased from 0 to 13mm), but not to the other antibiotics tested. The vancomycin sensitization was confirmed using MIC determinations (Table 1). Subsequent MIC determinations demonstrated that the intrinsic vancomycin resistance was reduced 8-fold for the Δ*spr* mutant and 32-fold for the Δ*spr*Δ*yebA* mutant (Table 1). Also, the Δ*spr*Δ*yebA* mutant revealed a moderate sensitization to novobiocin, penicillin G and rifampicin, while the sole Δ*yebA* mutant did not reveal any increase in sensitization to any of the antibiotics included, when compared to the wild type (Table 1).

**Table 1.**
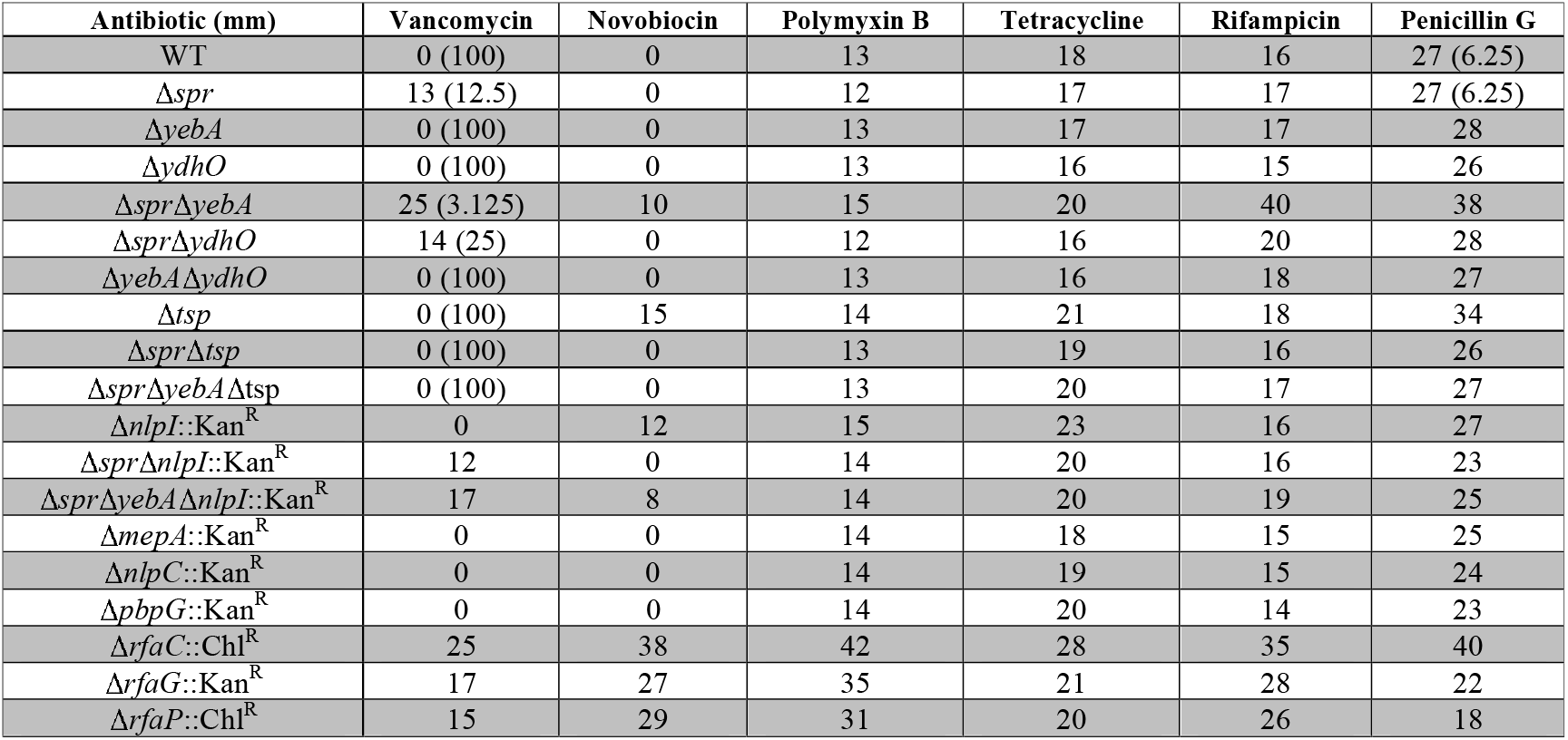
Antibiotic sensitivity profiles. Inhibition zones in millimeters (mm) on TY agar when antibiotics were applied on 6 mm diameter Whatman 3 filter paper discs. Numbers in parenthesis within the table are MIC in μg/ml determined in TY broth. Values are means of three independent replicates.

Any sensitization to the detergent deoxycholate (DOC) of the Δ*spr* and Δ*spr*Δ*yebA* mutants was evaluated by incubating them in 0.5% DOC for 30 minutes. In this assay only the Δ*spr*Δ*yebA* mutant revealed a clear sensitization to DOC exposure (Fig. 2A). These results implied that a plain Δ*spr* mutation in *S*. Typhimurium SR-11 associates with significant sensitization to vancomycin without revealing a general defect in the outer membrane permeability barrier, while addition of the Δ*yebA* mutation evidently caused an outer membrane destabilization.

**Figure 2.**
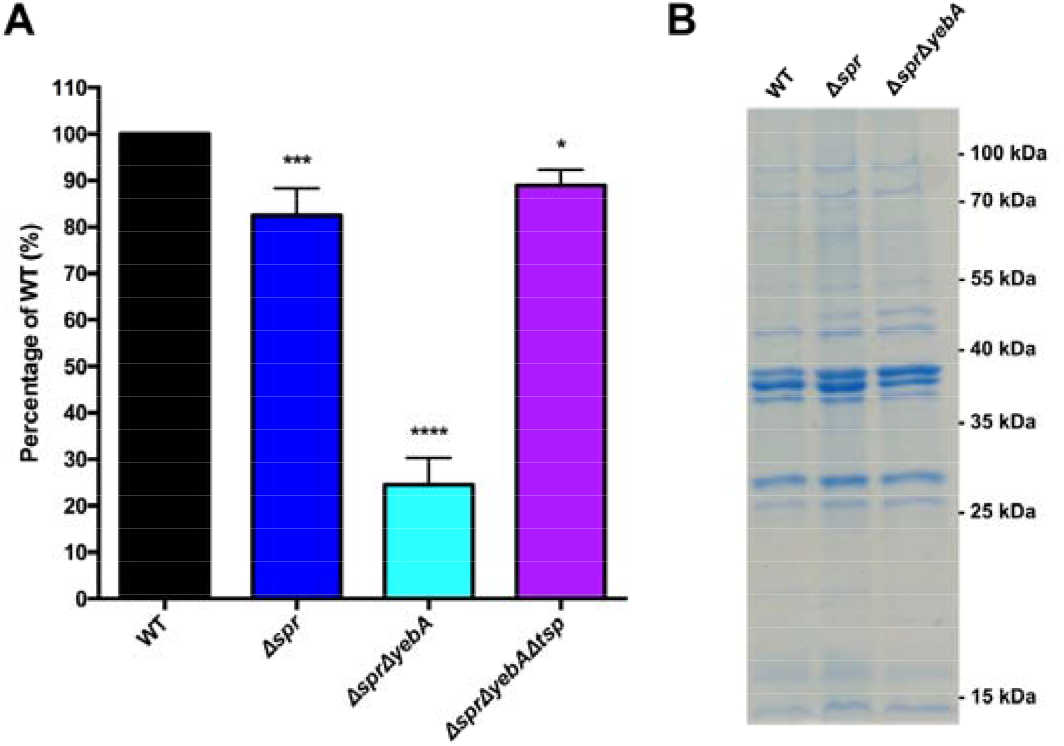
Detergent sensitivity and membrane protein profile of muramyl endopeptidase mutants. (**A**) For probing detergent sensitivity 10^5^ bacteria of each strains was incubated in 0.5% DOC for 30 min, where after the bacterial viable colonies were enumerated. Results are presented as relative CFU yield in relation to the wild-type. The graph shows the mean value and the standard error of the mean of three independent replicates. Statistical test was an one-way ANOVA with Dunnett’s correction for multiple comparisons when comparing the mutants to the wild type. (**B**) SDS-PAGE gel revealing the outer membrane protein profiles of the wild-type and mutant strains. The positions of prestained molecular weight markers are indicated at the right. The standard used is PageRuler Prestained Protein Ladder (#26616, Thermo Scientific).

### Selectivity of Spr in maintaining intrinsic vancomycin resistance

As the antibiotic sensitization profile caused by the Δ*spr* mutation was unexpected, we next tested whether this mutant phenotype was restricted to the SR-11 line of *S*. Typhimurium. Hence, the Δ*spr* mutation was introduced into the commonly used laboratory *S*. Typhimurium lines SL1344 and 14028. When tested for vancomycin tolerance, the MIC for the wild type SL1344 and 14028 strains was the same as for the wild type SR-11 line (100 μg/ml), whereas in the corresponding SL1344 and 14028 Δ*spr* mutant strains the MIC decreased to 12.5 μg/ml, equaling that of the SR-11 Δ*spr* mutant (Table 1).

In *E. coli* overproduction of PBP7 suppresses thermosensitive growth associated with a *mepS* mutation (Hiroshi Hara et al. 1996), suggesting that PBP7 and MepS connect in parallel pathways. Hence, we deleted *pbpG*, coding for the PBP7 homologue in *S*. Typhimurium. We also created deletion mutants for the *mepA* and *nlpC* homologues in *S*. Typhimurium, each coding for endopeptidases functionally related to Spr. Yet, none of the three additional mutants showed sensitization to the antibiotics included in the test panel (Table 1).

In *E. coli*, overexpression of selected outer membrane porin proteins can result in an outer membrane permeability defect (Krishnamoorthy et al. 2016). We compared outer membrane protein profiles of the wild type, Δ*spr* and Δ*spr*Δ*yebA* mutants on SDS-PAGE gels. We did not observe any significant difference in outer membrane protein profile for the mutants as compared to the wild type (Fig. 2B), arguing that the increased vancomycin sensitization is not caused by a major alteration in outer membrane protein composition.

Our observations combined show that deletion of *spr* in *S*. Typhimurium is associated with sensitization to vanco mycin, and that this sensitization is not restricted to line SR-11. Furthermore, we note that the lack of muramyl endopeptidases YebA, YdhO, PBP7, MepA or NlpC as such do not result in sensitization to vancomycin, nor does the *spr* deletion associate with a general overproduction of major porin proteins.

### Genetic verifications implicate a catalytically active Spr as the maintainer of intrinsic vancomycin tolerance in *S*. Typhimurium

To further ensure the PCR-based confirmations of the Δ*spr* and Δ*spr*Δ*yebA* mutations we performed whole genome sequencing on the two mutant constructs, which verified their expected genetic composition. To exclude any potential polar effects of the verified Δ*spr* deletions as a cause of vancomycin sensitization, we applied genetic complementation. We cloned the *spr* gene from *S*. Typhimurium SR-11 under the control of the arabinose-inducible promoter in the plasmid vector pBAD30. We next generated a point mutation in this plasmid replacing the codon for the conserved catalytic Spr cysteine residue with a codon for serine, creating a C70S alteration in the mature protein (Aramini et al. 2008; Singh et al. 2012). Introducing the cloned native *spr* gene into either the Δ*spr* or Δ*spr*Δ*yebA* mutant fully restored vancomycin resistance, whereas both the empty pBAD30 vector, and the plasmid coding for a catalytically inactive Spr variant, did not restore vancomycin resistance (Fig. 3). We conclude that intrinsic vancomycin resistance of S. Typhimurium requires a catalytically active Spr.

**Figure 3.**
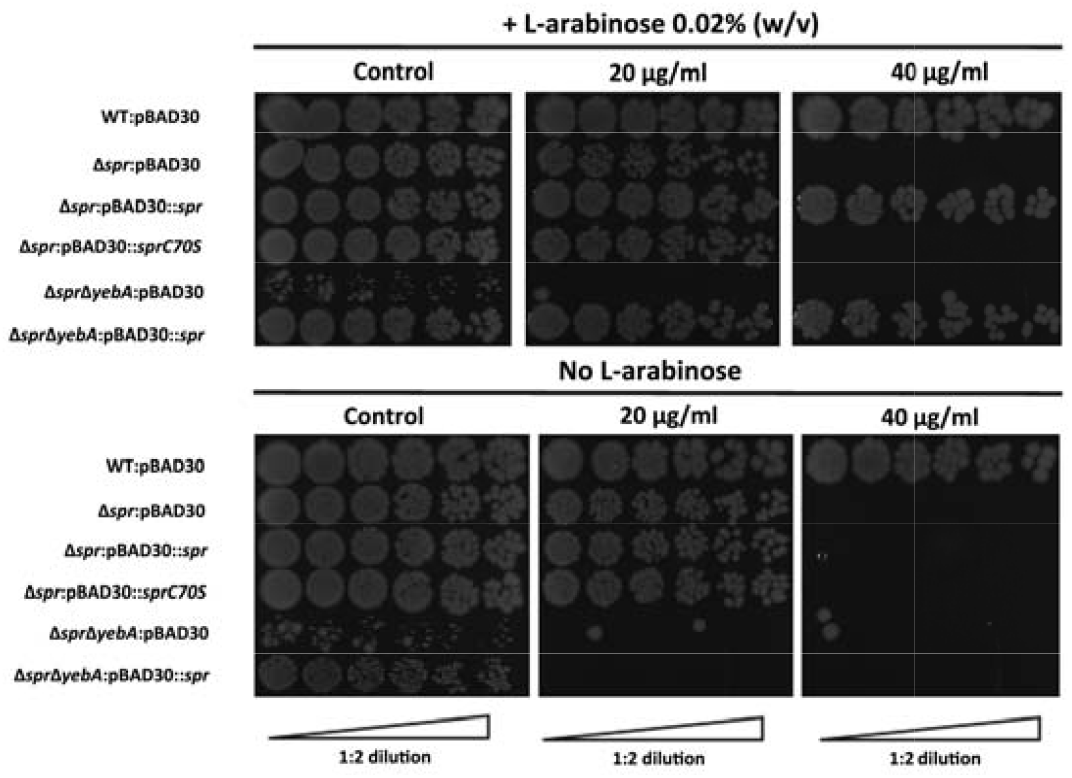
Drop-on-lawn visualization of complementation of vancomycin sensitivity in muramyl endopeptidase mutants. Wild-type, Δ*spr* and Δ*spr*Δ*yebA* mutants harboring either vector control pBAD30, pBAD30::spr for full complementation or pBAD30: *:sprC70S* as catalytic knockout were serially diluted and spread on TY agar plates, or TY agar plates supplemented with 20 μg/ml or 40 μg/ml of vancomycin, either containing L-arabinose or not. Images are representative of three replicates each.

### Effect of the Δ*spr* deletion on antibiotic-induced autolysis

Penicillin G activates in *E. coli* a protein-synthesis-dependent autolysis (Prestidge and Pardee 1957). Hence, we set out to test whether the Δ*spr* mutation would affect any autolytic behavior of *S*. Typhimurium in response to cell wall synthesis inhibitors. To enable quantification of bacterial cell lysis, we adapted a β–galactosidase (LacZ) release assays (detailed in Material and Methods). In this we incubated a logarithmic growth phase culture of *S*. Typhimurium with increasing concentrations of a cell wall synthesis inhibitor for 1 hour, after which the β–galactosidase activities were determined from the culture supernatants. Both the wild-type and Δ*spr* mutant revealed an increased level of β–galactosidase release as a function of increased concentration of vancomycin, with the release being more pronounced for the Δ*spr* mutant (Fig. 4A).

**Figure 4.**
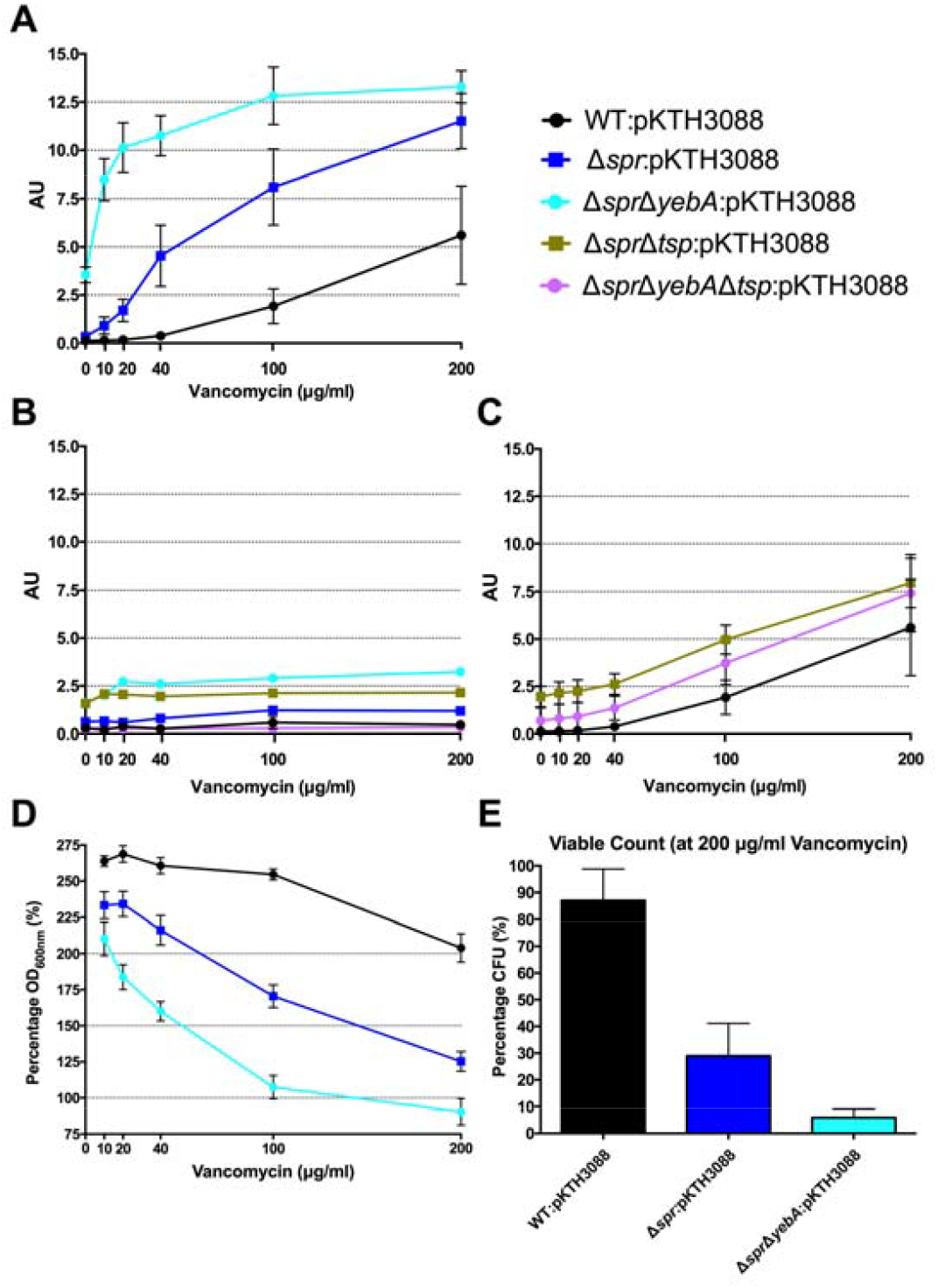
Vancomycin induced autolysis. Release of cytoplasmic β-galactosidase in the presence of increasing concentrations of vancomycin (**A & C**), or in combination with tetracycline (**B**) after 1 hour of incubation at 37°C. Decrease in optical density at 600nm with increasing vancomycin concentrations (**D**) and viable counts at highest concentration at 200 μg/ml vancomycin after one hour of incubation (**E**). Graph presents the mean values and standard error of the mean of three independent replicates.

To confirm that the vancomycin induced lysis depended on active protein synthesis we repeated the experiments with tetracycline added into the reaction mixture, as the wild-type and the Δ*spr* mutant exhibited identical MIC for tetracycline (Table 1). Adding tetracycline blocked the release of β–galactosidase from both strains (Fig. 4B).

As wild type *S*. Typhimurium SR-11 and the Δ*spr* mutant had an equal MIC for penicillin G (Table 3), we next repeated the lysis assay by replacing vancomycin with penicillin G. Both the wild-type and Δ*spr* mutant reached a similar plateau level of β–galactosidase release at a higher concentration range of penicillin G (Fig. 5A). However, compared to the wild-type, the β–galactosidase release was more pronounced for the Δ*spr* mutant at concentrations below the determined 6.25 μg/ml MIC for penicillin G (*t*-test: p < 0.01 for 2 and 4 μg/ml when comparing Δ*spr* mutant to wild-type). Yet again, as for vancomycin, the β–galactosidase release by penicillin G was blocked by addition of tetracycline (Fig. 5B).

**Figure 5.**
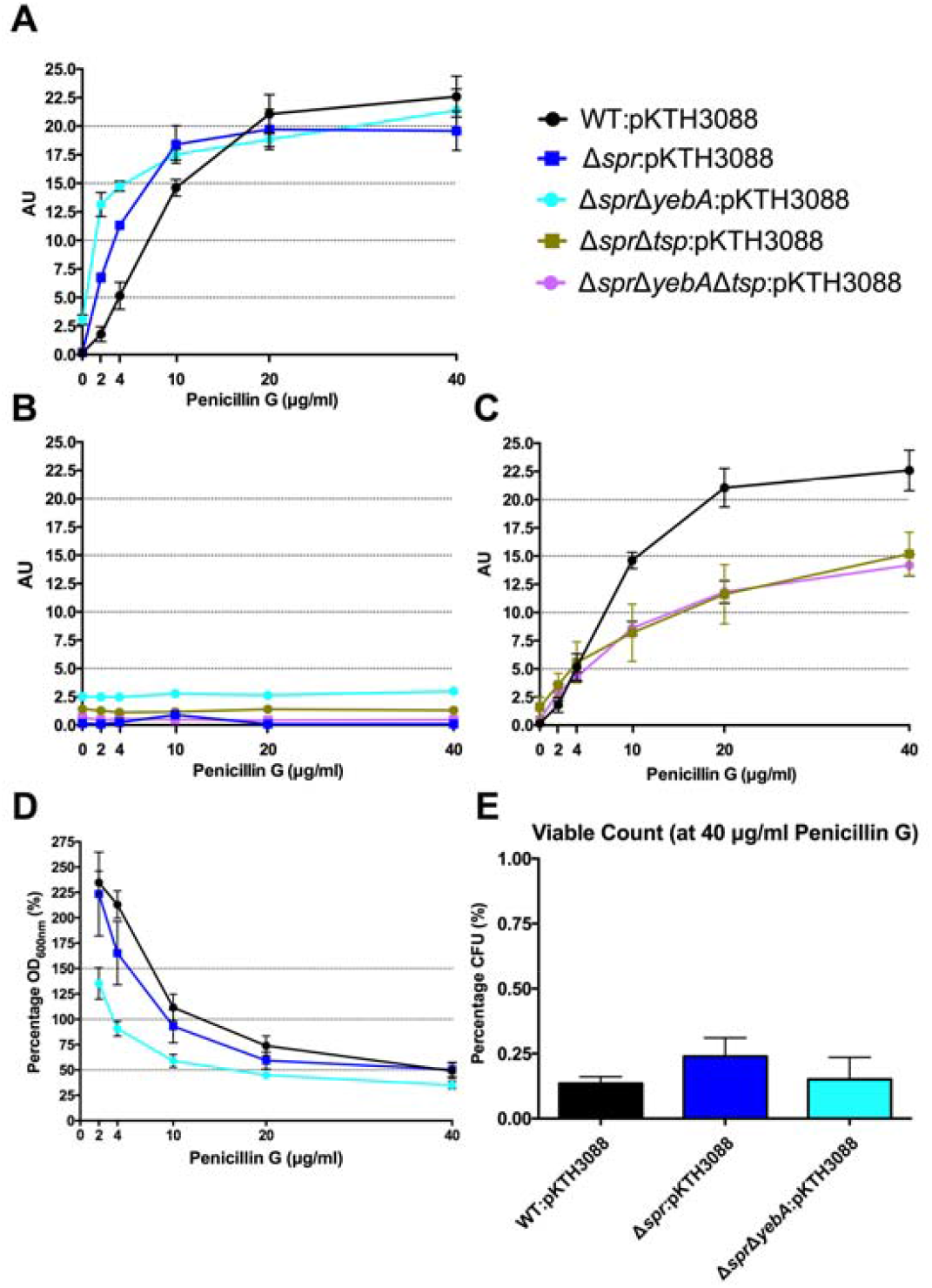
Penicillin G induced autolysis. Release of cytoplasmic β-galactosidase in the presence of increasing concentrations of penicillin G (**A & C**), or in combination with tetracycline (**B**) after 1 hour of incubation at 37°C. Decrease in optical density at 600nm with increasing penicillin G concentrations (**D**) and viable counts from highest concentration at 40 μg/ml penicillin G after one hour of incubarion (**E**). Graph presents the mean values and standard error of the mean of three independent replicates.

When we followed the development of the optical density (OD_60_0nm) under the 1 hour incubation with cell wall synthesis inhibitors (Fig. 4D and Fig. 5D), we noted that a substantial proportion of the bacteria apparently remained unlysed. When we determined the viable count from cultures exposed to 200 μg/ml of vancomycin, representing 2-fold MIC for wild-type and 8-fold MIC for Δ*spr* mutant bacteria, we recovered a substantial residual amount of viable bacteria from the cultures (Fig. 4E). When the viable counts were measured for the penicillin G exposed cultures (containing antibiotic 6 times the MIC), we could barely detect any viable bacteria (Fig. 5E).

The Δ*spr*Δ*yebA* mutant exhibited a substantial decrease in tolerance to both vancomycin and penicillin MIC as compared to either the wild-type or Δ*spr* mutant (Table 1). This increased sensitivity was associated with a significantly higher level of β–galactosidase release (relative to the wild-type or the Δ*spr* mutant) after exposure to either antibiotic for 1 hour.

Combined these data demonstrate that vancomycin induce a protein-synthesis-dependent autolysis in S. Typhimurium, and that intensity of this autolysis inversely correlated with the MIC to vancomycin. At the other hand, compared to the wild type, penicillin G evoked a more proficient autolysis in the Δ*spr* mutant despite the wild-type and Δ*spr* mutant had the same MIC for penicillin G.

### Isolation of Δ*spr* vancomycin-resistant suppressor mutations

Vancomycin-resistant mutants were selected in the Δ*spr*Δ*yebA* background using a vancomycin gradient plate (for details, see Materials and Methods). Twelve vancomycin-tolerant Δ*spr*Δ*yebA* mutants were analysed by whole genome sequencing. Eleven of the mutants carried overlapping deletions covering nucleotides 1,920,000 - 1,965,000 in the *S*. Typhimurium LT2 reference genome (Fig. 6A). At the centre of this region is *tsp*, encoding a periplasmic carboxypeptidase. The remaining suppressor mutant carried a point mutation within *tsp* itself (Fig. 6A). These data suggest that inactivation of *tsp* is the common feature of mutations that suppress the vancomycin sensitivity phenotype of the Δ*spr*Δ*yebA* mutant.

**Figure 6.**
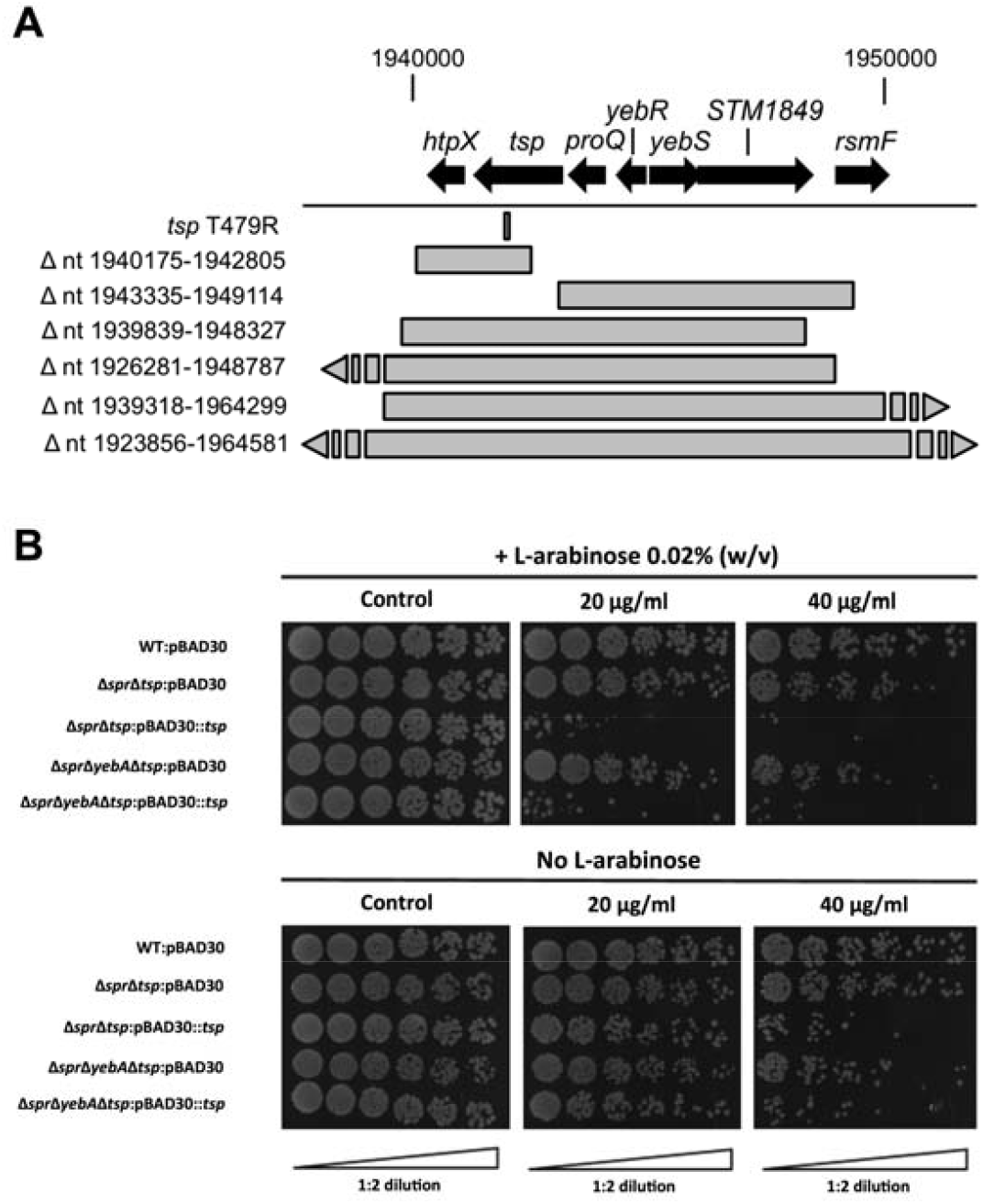
Suppression of vancomycin sensitivity in Δ*spr* mutant backgrounds. (**A**) Genomic visualization of vancomycin tolerant suppressor mutations with the bars representing genomic deletions, except for *tsp* T479R where the bar points to the position of a point mutation. (**B**) Wild-type and the Δ*tsp* suppressed mutants harboring either vector control pBAD30 or full complementation pBAD30::*tsp* were serially diluted and spread on TY agar plates, or TY agar plates supplemented with 20 μg/ml or 40 μg/ml of vancomycin, either containing L-arabinose or not. Images are representative of three replicates each.

To confirm the *tsp* mutations as suppressors, we deleted *tsp*, as well as the individual genes that mapped upstream and downstream of *tsp* (*proQ* and *htpX*) in the Δ*spr*Δ*yebA* mutant. Out of these three constructs only deletion of *tsp* resulted in a vancomycin resistant phenotype in the Δ*spr*Δ*yebA* mutant (Table 1). In addition, when the *tsp* deletion was introduced into the Δ*spr* mutant it converted the phenotype from vancomycin sensitive to vancomycin resistant (Table 1). Conversely, when a cloned *tsp* gene was introduced into the vancomycin resistant Δ*spr*Δ*tsp* and Δ*spr*Δ*yebA*Δ*tsp* mutants the phenotypes of each were reverted to vancomycin sensitive (Fig. 6B).

The Δ*tsp* mutation also reverted the general antibiotic sensitization of the Δ*spr*Δ*yebA* mutant (Table 1). In addition, introduction of the *tsp* deletion into the Δ*spr* and Δ*spr*Δ*yebA* mutants suppressed the autolysis by reducing their release of β–galactosidase in the presence of vancomycin or penicillin G (Fig. 4C and 5C). Thus, a *tsp* deletion acted as a general suppressor mutation for Δ*spr* and Δ*spr*Δ*yebA* mutant phenotypes. That said, the *tsp* deletion alone did not confer increased vancomycin resistance (Table 1).

In *E. coli*, Tsp co-purifies with the outer membrane lipoprotein NlpI, which assists Tsp in degrading MepS (Spr) (Singh et al. 2015). Furthermore, *nlpI* mutations suppress a temperature-sensitive phenotype of an *E. coli* Δ*mepS* mutant (Tadokoro et al. 2004) implying a functional connection between Tsp, NlpI and MepS. Yet, when we deleted *nlpI* in the *S*. Typhimurium Δ*spr* and Δ*spr*Δ*yebA* mutants, they retained their sensitization to vancomycin (Table 1). Hence the vancomycin-sensitive phenotype of the Δ*spr* mutant is dependent chiefly only on Tsp rather an on NlpI.

## DISCUSSION

In Gram-negative enteric bacteria, the outer membrane acts as a major barrier contributing to impermeability to large molecular compounds, adding to their intrinsic resistance to antibiotics, such as lysozyme, novobiocin, rifampicin and vancomycin (Grundström, Normark, and Magnusson 1980; Helander et al. 1989). Accordingly, alterations that disrupt the outer membrane barrier can sensitize enteric bacteria to such compounds (Sukupolvi et al. 1984). Lipopolysaccharide (LPS) strongly contributes to this outer membrane integrity and mutations in genes involved in LPS biosynthesis, including *envA*, *rfaC* (*waaC*), and *rfaP* (*waaP*), sensitize Gram-negative bacteria simultaneously to numerous antimicrobials (Grundström, Normark, and Magnusson 1980; Helander et al. 1989). Outer membrane integrity can also be disturbed by the expression of aberrant outer membrane proteins (Rhen, O’Connor, and Sukupolvi 1988), by exposing bacteria to polymyxin B nonapetide (Vaara and Vaara 1983; Ofek et al. 1994), a compound which lacks intrinsic bacteriocidal activity, or by a simultaneous genetic depletion of several murein hydrolases (Heidrich et al. 2002).

In *E. coli*, mutants lacking the Spr, YebA or YhdO single muramyl endopeptidases have no obvious growth phenotypes (Singh et al. 2012). In accordance, we observed that individual genetic deletion of murein endopeptidases Spr, YebA or YhdO in *S*. Typhimurium did not affect growth of the bacterium in broth cultures. We also created Δ*spr*, Δ*yebA* and Δ*yhdO* double mutants, to test for redundancy in their contribution to *S*. Typhimurium growth in broth. Specifically for the Δ*spr*Δ*yebA* mutant, we noted a minor defect in growth capacity. As this phenotype was not seen for the single Δ*spr*, Δ*yebA* or Δ*yhdO* mutants, or in the Δ*yebA*Δ*yhdO* double mutant, our observations suggested that *spr* might have a non-redundant function the growth conditions tested.

Previously outer membrane destabilization with accompanying vancomycin sensitivity has been seen in *E. coli* when deleting several muramyl hydrolases simultaneously (Korsak, Liebscher, and Vollmer 2005). Our study is the first where depletion of a single muramyl endopeptidase, Spr, results in vancomycin sensitization. Significantly, the vancomycin sensitization associated with the Δ*spr* mutation did not associate with sensitization to rifampicin or novobiocin, which would be expected in the case of classical outer membrane destabilization. This implies that there might be an alternative pathway for outer membrane leakage in an Spr-deficient strain.

This alternative pathway could involve an increased propensity for autolysis in bacteria lacking Spr, which itself could cause a form of cell wall destabilization. In support of this we could correlate increased autolytic behaviour upon exposure to vancomycin, measured as release of the cytoplasmic β–galactosidase enzyme. Using this assay, the Δ*spr* mutant also revealed a more rapid onset of autolytic behaviour upon exposure to penicillin G, despite the fact that the wild-type and Δ*spr* mutant shared the same MIC for penicillin G. Moreover, the Δ*spr* mutant appeared to undergo autolysis at sub-MIC concentrations of penicillin G.

When recording autolysis by measuring decrease in optical density, we noted that in the bacterial cultures undergoing lysis, exposed either to vancomycin or penicillin G, a significant proportion of each culture remained apparently non-lysed. Also, even at a vancomycin concentration 16-fold MIC, we could recover a substantial proportion of viable Δ*spr* bacteria at the end of the autolysis experiment. Viable Δ*spr* bacteria were also recovered from the corresponding penicillin G-exposed cultures but at a 100-fold lower frequency. These results would imply that vancomycin and penicillin G both induce autolysis in the Δ*spr* mutant, but that in each case there remains a sub-population that is resilient to autolysis, with a substantial proportion of cultivable bacteria remaining 1 hour after vancomycin exposure that exceeds their MIC value. An hypothesis to explain these data is that there is a different proportions of lysis-proficient and non-proficient bacteria in Δ*spr* mutant and wild-type bacterial cultures. Accordingly, the effect of deleting *tsp* in the dynamics of autolysis could be that it decreases the proportion of the lysis-proficient bacteria. Whether these non-proficient bacteria would eventually lose their colony-forming ability, or lyse, upon prolonged vancomycin exposure remains to be evaluated.

In Gram-positive bacteria vancomycin resistance is achieved through the acquisition of large genetic blocks coding for new peptidoglycan motifs, rather than through point mutations (Faron, Ledeboer, and Buchan 2016; Gardete and Tomasz 2014). As the MIC for our vancomycin-sensitive Δ*spr*Δ*yebA* mutant approached MIC values of susceptible Gram-positive pathogens such as *Enterococcus faecalis* and *Staphylococcus aureus* (Susceptible Enterococci spp. ≤ 4 μg/ml and susceptible *S. aureus* ≤ 2 μg/ml according to “EUCAST: Clinical Breakpoints” (http://www.eucast.org)), we set out to probe whether we could in a Gram-negative vancomycin-sensitive bacterium select for spontaneous mutations adding to vancomycin tolerance. In this, we selected and genetically confirmed that Δ*spr*-mediated vancomycin sensitization, even the more pronounced sensitization of the Δ*spr*Δ*yebA* mutant, required a *tsp*-proficient genetic background. The gene *tsp* itself codes for periplasmic endopeptidase implicated in processing of Spr and PBP3 (Singh et al. 2015; H Hara et al. 1991; H. Hara et al. 1989). While any corresponding Spr/Tsp machinery does not exist in enterococci, the success of isolating vancomycin resistant mutations, even point mutations, in our Δ*spr*Δ*yebA* mutant background adds to our understanding of how vancomycin is involved in blocking peptidoglycan synthesis by showing that resistance can be achieved without horizontal gene transfer. We here also demonstrate that intrinsic *S*. Typhimurium vancomycin resistance relies on a catalytically active Spr. This, and the fact that the vancomycin sensitivities of the Δ*spr* and Δ*spr*Δ*yebA* mutants are close to the clinical breakpoints of relevant clinical Gram-positive pathogens places the catalytic activity of Spr as a potential target whose inhibition could potentiate treatment of enteric bacterial infections with vancomycin.

In summary, we have genetically defined a new pathway for intrinsic resistance to the large-molecular-weight antibiotic vancomycin, that is not dependent on outer membrane permeability, in the Gram-negative pathogen *S*. Typhimurium. This new pathway involves the combined action of the muramyl endopeptidases Spr, together with the protease Tsp, in maintaining the peptidoglycan homeostasis essential for maintaining the cell wall integrity of the bacterium upon antibiotic challenge. These insights add to the knowledge needed to combat the increasing problem of antibiotic resistance in Gram-negative bacteria.

## MATERIALS AND METHODS

### Bacterial strains

Mutations were constructed in the *Salmonella enterica* serovar Typhimurium SR-11 background (Sukupolvi et al. 1997), and are listed in Table S1. The strains LB5010 (Bullas and Ryu 1983) and ATCC 14028 were used as intermediary hosts in creating mutations. *Escherichia coli* Top10 and TG1 were used as intermediary hosts during cloning. Furthermore, S. Typhimurium strains ATCC 14028s and SL1344 were also used to host a Δ*spr* mutation.

### Media and growth conditions

Growth media included tryptone and yeast extract (Sigma-Aldrich, 10 g/l respective 5 g/l) with 10 g/l of NaCl (LB medium), or without NaCl (TY medium). Cultures were incubated at 37°C unless otherwise stated. When needed, antibiotics were added to the growth media at the following concentrations: ampicillin 100 μg/ml; chloramphenicol 25 μg/ml; kanamycin 50 μg/ml; tetracycline 10 μg/ml. All antibiotics were purchased from Sigma-Aldrich (Sweden).

For determining growth curves, bacteria were incubated overnight in 2ml LB at 220 rpm in 37°C. The next, day 150μl of the culture was mixed with 850μl of PBS and the OD_600nm_ measured. The bacteria were then adjusted to an OD_600nm_ of 0.25 (Ultrospec 1000, Pharmacia Biotech). From this OD_600nm_ the bacteria were then diluted 1:25 in either LB or TY broth resulting in a final OD_600nm_ of 0.01. 400μl of this bacterial suspension was then loaded into wells in a Honeycomb Bioscreen plate (OY Growth Curves AB LTd, Helsinki, Finland) in three technical replicates. Uninoculated media was used as a negative growth control. The Bioscreen C plate reader (OY Growth Curves AB Ltd) was set to an OD of 600nm and optical density measurement was taken every 15 minutes with 5 seconds of agitation before every measurement, up until 24 hours.

### PCR and oligonucleotides

Polymerase chain reaction (PCR) was performed using an Eppendorf Mastercycler Personal. Oligonucleotides were designed using the genome of *S*. Typhimurium LT2 as reference. For the generation of the inserts for gene deletions the PCR was performed using Phusion High-Fidelitiy PCR master mix with HF buffer (New England Biolabs, USA). The cycling conditions were as following: 98°C for 1 min and 30 cycles of 98°C for 15 seconds, 54.5°C for 10 seconds, 72°C for 40 seconds, and 72°C for 2 minutes. The oligonucleotides were ordered from Sigma-Aldrich and specified in Table S2.

For routine confirmatory PCR, Phusion High-Fidelitiy PCR master mix with HF buffer was used. The cycling conditions were as follows: 98°C for 1 minute and 30 cycles of 98°C for 15 seconds, annealing temperature for 10 seconds, 72°C for elongation time, and 72°C for 2 minutes. Annealing temperature varied depending on the primer pairs used, and elongation time was based on the length of the expected product (30 seconds per kilobase). Oligonucleotide sequences are shown in Table S2.

### Bacteriophage transduction

Phage P22*int* was that of Schmieger (Schmieger 1972). Transducing phages were prepared on strain LB5010 (in LB broth supplemented with D-galactose (Fluka BioChemika) to 0.2% (wt/vol)) or strain 14028s carrying the mutation of interest. The next day chloroform was added to the culture and the culture was vortexed. The culture was then centrifuged for 10 minutes at 18500 g to create phase separation. The top phase was recovered and used to transfer the genetic marker. The transduction into *S*. Typhimurium SR-11 was done by incubating 20μl of the P22*int* phage containing the genetic marker with 1ml of exponential phase culture. These were incubated at 37°C with shaking at 220 rpm for 1 hour and after washing in PBS plated onto appropriate selective LB agar plates.

### Construction and isolation of mutants

The antibiotic marker amplified from either pKD3 (Chl^R^) or pKD4 (Kan^R^) was conducted using primers with 3’-ends overlapping the borders of the gene to be deleted, and subsequently inserted into the chromosome, to replace the gene of interest, by double-stranded DNA lambda-red recombineering (Datsenko and Wanner 2000; Yu et al. 2000). As recipients we used *S*. Typhimurium strain LB5010 containing the pSIM6, or *S*. Typhimurium strain 14028 containing pSIM5-tet plasmid, each grown to an OD_600nm_ of approximately 0.3 at 32°C with shaking at 220 rpm in a water bath. To induce the lambda-red genes the bacteria were transferred to a 42°C water bath shaking at 220 rpm for 15 minutes. After cooling on ice for 10 minutes, bacterial cells were made electrocompetent by washing with ice-cold deionized water four times.

Electroporation of the PCR products generated from pKD3 or pKD4 was done using a Gene Pulser (Bio-Rad, USA) by mixing 25μl of electrocompetent cells and 0.5μl of purified PCR product, with settings 1.8 kV, 25 F and 200Ω. Cells were recovered in 1ml of LB at 32°C and 220 rpm for 2 hours. After recovery the culture was spun down and the pellet was spread on LB agar plates containing either chloramphenicol or kanamycin to select for recombinants. The genetic marker was subsequently transferred to wild type *S*. Typhimurium SR-11 by phage P22*int* transduction from either the LB5010 or 14028 mutant strains.

Antibiotic markers were removed from the mutants using the plasmid pCP20. Briefly, pCP20 was transferred by P22*int* transduction to the recipient strain containing the antibiotic marker, with selection for colonies on LB agar plates containing ampicillin. Transductants were subcultured three times at 28°C on selective LB agar plates. The bacteria were then subcultured three more times at 37°C to select for loss of plasmid and loss of antibiotic marker. The loss of the antibiotic marker was confirmed by a diagnostic PCR.

Isolation of vancomycin-tolerant Δ*spr*Δ*yebA* mutants was conducted by spreading a LB broth culture of the Δ*spr*Δ*yebA* mutant on a vancomycin gradient TY agar plate. In this, a plate of 14 cm in diameter was poured in a tilted position with 37.5ml of TY agar containing vancomycin at 40 μg/ml. After solidification, 37.5ml TY agar lacking antibiotic was poured on top of the solidified TY agar containing vancomycin, now in a horizontal plane. The plate was seeded with about 10^7^ CFU of Δ*spr*Δ*yebA* mutant bacteria in their logarithmic growth phase. After 16 hours of incubation yielded colonies were isolated at the higher concentration end of the gradient.

### Plasmid constructions and site-directed mutagenesis

For creating plasmids for genetic complementation, *spr* and *tsp* were PCR amplified using S. Typhimurium SR-11 genomic DNA as template, using oligonucleotide primers to create suitable restriction sites at each end of the amplified fragment. Enzymes used for restriction digestion were SacI or EcoRI, and HindIII (New England Biolabs) while T4 DNA ligase was used to ligate *spr* fragments into vector pBAD30 (Guzman et al. 1995). Reaction buffers were provided by the enzyme supplier. Following ligation, plasmids were transformed (as above) into chemically competent *E.coli* TG1 or Top10 cells (Invitrogen) from where the constructs were purified and electroporated (as above) into *S*. Typhimurium LB5010. The plasmids were then transferred into *S*. Typhimurium SR-11 by *P22int* transduction.

### Whole-genome sequencing and analysis

Genomic DNA was prepared from bacterial cultures using a Masterpure DNA Purification Kit (Epicentre). Libraries for sequencing were prepared using Nextera XT sample preparation and index kits (Illumina). The quality of libraries was assessed using a Tapestation 2200 (Agilent) with high-sensitivity D5000 Screentape. Sequencing of the libraries was done using a Miseq device (Illumina) using a 600-cycle V3 reagent kit. The sequences were processed and analyzed using CLC Genomics Workbench 9.0.1 (CLC bio).

### Deoxycholate (DOC) sensitivity test

To assay detergent sensitivity, bacteria were diluted 1:100 in LB from an overnight culture, and grown in LB for 2 hours at 37°C with shaking at 220 rpm. The OD_600nm_ of the culture was measured, and the formula ((0.484/OD_600nm_) x 2.1) used to adjust the amount of bacteria to obtain approximately 10^7^ CFU/ml. The bacterial suspension was then diluted 1:100 in distilled water with sodium deoxycholate (DOC; Sigma-Aldrich, Sweden) freshly added to a final concentration of 0.5% (wt/vol). After an incubation of 30 minutes at room temperature, viable counts were determined from the DOC-suspension by plating dilutions on LB agar plates.

### SDS-PAGE gel electrophoresis

Polyacrylamide-bis-acrylamide gel electrophoresis was conducted according to Laemmli (Laemmli 1970), using custom made a 12.5% separation (pH 8.8) and a 6% stacking gel (pH 6.8). (Thermo Scientific, Sweden). For solubilisation, samples were suspended in reducing protein sample buffer (0.125 M Tris-HCl pH 6.5, 3.6% SDS, 10% β-mercaptoethanol, 2% glycerol, and bromphenol blue enough to give a deep blue color) and heated 10 minutes at 97°C before application on the gel. After completed electrophoresis, gels were stained using Imperial stain.

### Bacterial membrane fractionation

The outer membrane fraction was isolated according to Sabet and Schnaitman (Sabet and Schnaitman 1973). Bacteria from overnight LB cultures were diluted 1:100 into TY broth and grown at 37°C for 2 hours to mid exponential phase. The OD_600nm_ of the bacterial cultures were measured and then normalized. Bacteria were pelleted by centrifugation at 6000 g for 10 minutes, resuspended in PBS, cooled on ice, and disrupted by equal numbers of 10 seconds sonication pulses until the suspensions visibly cleared. Bacteria were removed by low-speed centrifugation (1500 g), after which the membrane fraction was pelleted by high-speed centrifugation (18500 g, 10 minutes). The waxy brownish pellet was then re-suspended in 50mM Tris-HCl buffer containing 10mM MgCl_2_ and 0.5% Triton X-100. The membrane fraction was again pelleted by centrifugation (18500 g, 10 minutes), and the re-suspension and high-speed centrifugation steps were repeated. The final colourless pellet was then suspended in reducing protein sample buffer and run on a 12.5% according to Laemmli (Laemmli 1970) following by staining with Imperial stain (Thermo Scientific).

### Disk diffusion sensitivity testing

Bacteria were grown overnight in LB broth at 37°C and 220 rpm. The overnight culture was diluted 1:100 in LB broth and grown for 2 hours at 37°C and 220 rpm. The OD_600nm_ was measured of the 2 hour culture and the formula ((0.484/OD_600nm_) x 2.1) was used to calculate the amount of bacteria needed for 10^6^ CFU/ml. 3 x 10^5^ CFU/ml bacteria were then spread on large (13.7 cm diameter) TY agar plates and the antibiotic disks were placed on top of the bacteria and the plates were incubated overnight at 37°C. The next day the diameters of the inhibitory zones were measured.

The disks, 6mm in diameter, were custom-made out of Whatman 3 paper, using an ordinary office paper puncher. Each disk then was infused with 5μl of an antibiotic, and then let to dry. Concentrations of the antibiotics used were; tetracycline 10 μg/ml, vancomycin 20 μg/ml, rifampicin 10 μg/ml, polymyxin B 10 μg/ml, novobiocin 10 μg/ml, and penicillin G 10 μg/ml (Sigma-Aldrich, Sweden). Every antibiotic was dissolved in water except rifampicin, which was dissolved in DMSO.

### MIC determination

Bacteria were grown overnight in LB broth at 37°C and 220 rpm. The overnight culture was diluted 1:100 in LB broth and grown for 2 hours at 37°C and 220 rpm. The OD_600nm_ was measured of the 2 hour culture and the formula ((0.484/OD_600nm_) x 2.1) was used to calculate the amount of bacteria needed for 2 x 10^4^ CFU/ml in TY broth. This dilution was subsequently pipetted into a 96-well plate, containing TY broth with either vancomycin or penicillin G, resulting in a final concentration of 10^4^ CFU/ml bacteria. The highest concentration for the vancomycin MIC testing was 100 μg/ml and 50 μg/ml for penicillin G. Then, the antibiotic was diluted down in steps of 1:2 from the highest concentration before an incubation over night at 37°C.

### Drop-on-lawn

Bacteria were grown overnight in LB broth at 37°C and 220 rpm. The overnight culture was diluted 1:100 in LB broth and grown for 2 hours at 37°C and 220 rpm. The OD_600nm_ was measured of the 2 hour culture and the formula ((0.484/OD_600nm_) x 2.1) was used to calculate the amount of bacteria needed for 10^6^ CFU/ml. From 10^6^ CFU/ml a 1:2 serial dilution series was made into 1ml PBS. From each dilution a 5μl droplet was pipetted onto TY agar plates containing none, 20 μg/ml, or 40 μg/ml vancomycin and the TY agar plates were incubated overnight at 37°C. For genetic complementation tests TY agar plates were supplemented with 0.02% (weight/vol) L-arabinose (Sigma-Aldrich, Sweden).

### β-galactosidase (LacZ) release assay

A phage lysate was prepared from LB5010 carrying pKTH3088 (Taira et al. 1991), which is a pACYC184 derivative expressing the *E.coli lacZ* gene producing β-galactosidase (Casadaban et al. 1983). This P22 lysate was used transfer the pKTH3088 to *S*. Typhimurium SR-11 strains. To assay for the release of β-galactosidase (LacZ), bacteria were grown overnight in LB broth at 37°C and 220 rpm. The overnight culture was diluted 1:100 in TY broth and grown for 2 hours at 37°C and 220 rpm to mid exponential phase. Following the incubation the OD_600nm_ of the culture was measured, where after 300μl of bacteria was added to 700μl TY broth containing different concentrations of penicillin G or vancomycin and incubated at 37°C for 60 minutes. When attempting to inhibit penicillin G and vancomycin induced autolysis, 20 μg/ml tetracycline was added in this step. The bacteria were then pelleted by centrifuging and 200 μl of the supernatant was transferred into a reaction tube containing 600μl reaction buffer (0.001M MgSO_4_ and 0.05M β-mercaptoethanol (Sigma-Aldrich, Sweden) in 0.01M PBS, pH 7.2), and 200μl of 4 mg/ml ONPG (Sigma-Aldrich, Sweden) dissolved in reaction buffer. The β-galactosidase activity was stopped at given time points by adding 500μl of 0.5M sodium carbonate. 1ml of the samples were transferred into cuvettes and the β-galactosidase activity was measured using OD_420nm_. The formula to calculate the arbitrary units (AU) was implemented as according to Miller (Miller 1972).

When measuring for the proportional and total β-galactosidase activities in the bacterial cells the samples were similarly incubated overnight and diluted 1:100 the next day for the 2 hour incubation. After the incubation the OD_600nm_ was measured and 300μl was added to 700μl of TY broth containing 40μg/ml penicillin G and incubated for 60 minutes at 37°C. The samples were pelleted by centrifugation and 200μl of the supernatant was added to 600μl of Z-buffer and 200μl of 4mg/ml ONPG in reaction buffer. In order to assay the amount of LacZ in the pellet, the rest of the supernatant was discarded and the pellet resuspended in 100μl TY broth. This suspension was added to 600μl of reaction buffer and 200μl of 4 mg/ml ONPG in reaction buffer supplemented with 50μl of 0.1% SDS (Sigma-Aldrich, Sweden) and 50μl of >99% chloroform (Sigma-Aldrich, Sweden) to allow for the ONPG to penetrate into the pelleted cells. The samples were then incubated at 30°C for 20 minutes, sodium carbonate was added, the OD_420nm_ was measured, and the calculations for the arbitrary units performed as above.

### OD_600nm_ determination following antibiotic treatment

As a complement to the β-galactosidase release assay we followed alterations in OD_600nm_ for the same strains. The strains were grown as above in that the bacteria were grown overnight in LB broth at 37°C and 220 rpm. The overnight culture was diluted 1:100 in TY broth and grown for 2 hours at 37°C and 220 rpm to mid exponential phase. Following the incubation the OD_600nm_ of the culture was measured as the “pre-value”. Simultaneously, 300μl of bacteria was added to 700μl TY broth containing different concentrations of vancomycin or penicillin G and incubated at 37°C for 60 minutes. Following this incubation the OD_600nm_ was measured as the “post-value”. To visualize the effect of the antibiotics on the OD_600nm_ the post-values were divided with the pre-values.

### Viable count from broth

Also as a complement to the β-galactosidase release assays we determined the viable count for each strain at the highest concentration of vancomycin (200 μg/ml) and penicillin G (40 μg/ml). The strains were grown as above in that the bacteria were grown overnight in LB broth at 37°C and 220 rpm. The overnight culture was diluted 1:100 in TY broth and grown for 2 hours at 37°C and 220 rpm to mid exponential phase. Following the incubation the viable count for the input was enumerated by taking 300μl of bacteria into 700μl TY broth and a 1:10 serial dilution was performed in PBS. Bacteria were then spread on LB agar plates and incubated overnight at 37°C and the CFU were counted the next day, yielding the input value. In parallel 300μl of bacteria was added to 700μl TY broth containing either penicillin G or vancomycin and incubated at 37°C for 60 minutes. Following this incubation the strains were serially diluted 1:10 in PBS and the bacteria were then spread on LB agar plates and incubated overnight at 37°C and the CFU was counted the next day, yielding the output value.

### Statistical analysis

GraphPad Prism v6.0g (GraphPad Software, Inc., USA) was used for statistical analysis.

## ACKNOWLEDGEMENTS

We thank Sandra Muschiol for making electrocompetent *E. coli* Top10 cells, and Edmund Loh for giving input on the manuscript.

## FUNDING

This work was supported by Vetenskapsrådet (the Swedish Research Council) grants Dnr 4-30 16-2013 (M.R.), 2013-8643 (D.H.), 2016-04449 (D.H.), and 2017-03953 (D.H.). M.R. is also supported as a visiting professor in the Umeå Centre for Microbial Research (UCMR) Linnaeus Program by grant Dnr 349-2007-8673 from Vetenskapsrådet.

## AUTHOR CONTRIBUTION STATEMENT

KV, HW, DH, and MR designed the study. KV, DLH, IS, and MR performed experiments. KV, DH, and MR wrote the manuscript.

## CONFLICT OF INTEREST STATEMENT

Author have no conflicts of interest. The funders had no role in study design, data collection and interpretation, or the decision to submit the work for publication.

## REFERENCES

Anderle, Christine, Martin Stieger, Matthew Burrell, Stefan Reinelt, Anthony Maxwell, Malcolm Page, and Lutz Heide. 2008. “Biological Activities of Novel Gyrase Inhibitors of the Aminocoumarin Class.” Antimicrobial Agents and Chemotherapy 52 (6): 1982–90. https://doi.org/10.1128/AAC.01235-07.

Angelo, Kristina M., Jared Reynolds, Beth E. Karp, Robert Michael Hoekstra, Christina M. Scheel, and Cindy Friedman. 2016. “Antimicrobial Resistance Among Nontyphoidal Salmonella Isolated From Blood in the United States, 2003–2013.” The Journal of Infectious Diseases 214 (10): 1565–70. https://doi.org/10.1093/infdis/jiw415.

Aramini, James M., Paolo Rossi, Yuanpeng J. Huang, Li Zhao, Mei Jiang, Melissa Maglaqui, Rong Xiao, et al. 2008. “Solution NMR Structure of the NlpC/P60 Domain of Lipoprotein Spr from Escherichia Coli: Structural Evidence for a Novel Cysteine Peptidase Catalytic Triad.” Biochemistry 47 (37): 9715–17. https://doi.org/10.1021/bi8010779.

Baquero, M. R., M. Bouzon, J. C. Quintela, J. A. Ayala, and F. Moreno. 1996. “dacD, an Escherichia Coli Gene Encoding a Novel Penicillin-Binding Protein (PBP6b) with DD-Carboxypeptidase Activity.” Journal of Bacteriology 178 (24): 7106–11.

Beggs, William H., and Fred A. Andrews. 1976. “Role of Ionic Strength in Salt Antagonism of Aminoglycoside Action on Escherichia Coli and Pseudomonas Aeruginosa.” The Journal of Infectious Diseases 134 (5): 500–504. https://doi.org/10.1093/infdis/134.5.500.

Botta, G. A., and J. T. Park. 1981. “Evidence for Involvement of Penicillin-Binding Protein 3 in Murein Synthesis during Septation but Not during Cell Elongation.” Journal of Bacteriology 145 (1): 333–40.

Broome-Smith, J. K., and B. G. Spratt. 1982. “Deletion of the Penicillin-Binding Protein 6 Gene of Escherichia Coli.” Journal of Bacteriology 152 (2): 904–6.

Bullas, L R, and J I Ryu. 1983. “Salmonella Typhimurium LT2 Strains Which Are R-M+ for All Three Chromosomally Located Systems of DNA Restriction and Modification.” Journal of Bacteriology 156 (1): 471–74.

Casadaban, M. J., A. Martinez-Arias, S. K. Shapira, and J. Chou. 1983. “Beta-Galactosidase Gene Fusions for Analyzing Gene Expression in Escherichia Coli and Yeast.” Methods in Enzymology 100: 293–308.

Coque, T. M., F. Baquero, and R. Canton. 2008. “Increasing Prevalence of ESBL-Producing Enterobacteriaceae in Europe.” Euro Surveillance: Bulletin Europeen Sur Les Maladies Transmissibles = European Communicable Disease Bulletin 13 (47).

Datsenko, K A, and B L Wanner. 2000. “One-Step Inactivation of Chromosomal Genes in Escherichia Coli K-12 Using PCR Products.” Proceedings of the National Academy of Sciences of the United States of America 97 (12): 6640–45. https://doi.org/10.1073/pnas.120163297.

Delcour, Anne H. 2009. “Outer Membrane Permeability and Antibiotic Resistance.” Biochimica et Biophysica Acta (BBA) - Proteins and Proteomics, Mechanisms of Drug Efflux and Strategies to Combat Them, 1794 (5): 808–16. https://doi.org/10.1016/j.bbapap.2008.11.005.

Denome, S. A., P. K. Elf, T. A. Henderson, D. E. Nelson, and K. D. Young. 1999. “Escherichia Coli Mutants Lacking All Possible Combinations of Eight Penicillin Binding Proteins: Viability, Characteristics, and Implications for Peptidoglycan Synthesis.” Journal of Bacteriology 181 (13): 3981–93.

“EUCAST: Clinical Breakpoints.” n.d. Accessed July 6, 2018. http://www.eucast.org/clinical_breakpoints/.

Faron, Matthew L., Nathan A. Ledeboer, and Blake W. Buchan. 2016. “Resistance Mechanisms, Epidemiology, and Approaches to Screening for Vancomycin-Resistant Enterococcus in the Health Care Setting.” Journal of Clinical Microbiology 54 (10): 2436–47. https://doi.org/10.1128/JCM.00211-16.

Gardete, Susana, and Alexander Tomasz. 2014. “Mechanisms of Vancomycin Resistance in Staphylococcus Aureus.” The Journal of Clinical Investigation 124 (7): 2836–40. https://doi.org/10.1172/JCI68834.

Gray, Andrew N., Alexander JF Egan, Inge L. van’t Veer, Jolanda Verheul, Alexandre Colavin, Alexandra Koumoutsi, Jacob Biboy, et al. 2015. “Coordination of Peptidoglycan Synthesis and Outer Membrane Constriction during Escherichia Coli Cell Division.” eLife 4 (May): e07118. https://doi.org/10.7554/eLife.07118.

Grundström, T., S. Normark, and K. E. Magnusson. 1980. “Overproduction of Outer Membrane Protein Suppresses envA-Induced Hyperpermeability.” Journal of Bacteriology 144 (3): 884–90.

Guzman, L. M., D. Belin, M. J. Carson, and J. Beckwith. 1995. “Tight Regulation, Modulation, and High-Level Expression by Vectors Containing the Arabinose PBAD Promoter.” Journal of Bacteriology 177 (14): 4121–30.

Hara, Hiroshi, Noriko Abe, Masayo Nakakouji, Yukinobu Nishimura, and Kensuke Horiuchi. 1996. “Overproduction of Penicillin-Binding Protein 7 Suppresses Thermosensitive Growth Defect at Low Osmolarity due to an Spr Mutation of Escherichia Coli.” Microbial Drug Resistance 2 (1): 63–72. https://doi.org/10.1089/mdr.1996.2.63.

Hara, H., Y. Nishimura, J. Kato, H. Suzuki, H. Nagasawa, A. Suzuki, and Y. Hirota. 1989. “Genetic Analyses of Processing Involving C-Terminal Cleavage in Penicillin-Binding Protein 3 of Escherichia Coli.” Journal of Bacteriology 171 (11): 5882–89.

Hara, H, Y Yamamoto, A Higashitani, H Suzuki, and Y Nishimura. 1991. “Cloning, Mapping, and Characterization of the Escherichia Coli Prc Gene, Which Is Involved in C-Terminal Processing of Penicillin-Binding Protein 3.” Journal of Bacteriology 173 (15): 4799–4813.

Harder, Karen J., Hiroshi Nikaido, and Michio Matsuhashi. 1981. “Mutants of Escherichia Coli That Are Resistant to Certain Beta-Lactam Compounds Lack the ompF Porin.” Antimicrobial Agents and Chemotherapy 20 (4): 549–52.

Heidrich, Christoph, Astrid Ursinus, Jürgen Berger, Heinz Schwarz, and Joachim-Volker Höltje. 2002. “Effects of Multiple Deletions of Murein Hydrolases on Viability, Septum Cleavage, and Sensitivity to Large Toxic Molecules in Escherichia Coli.” Journal of Bacteriology 184 (22): 6093–99. https://doi.org/10.1128/JB.184.22.6093-6099.2002.

Heijenoort, Jean van. 2011. “Peptidoglycan Hydrolases of Escherichia Coli.” Microbiology and Molecular Biology Reviews: MMBR 75 (4): 636–63. https://doi.org/10.1128/MMBR.00022-11.

Helander, Ilkka M., Martti Vaara, Soila Sukupolvi, Mikael Rhen, Sirkku Saarela, Ulrich Zähringer, and P. Helena Mäkelä. 1989. “rfaP Mutants of Salmonella Typhimurium.” European Journal of Biochemistry 185 (3): 541–46. https://doi.org/10.1111/j.1432-1033.1989.tb15147.x.

Henderson, T. A., M. Templin, and K. D. Young. 1995. “Identification and Cloning of the Gene Encoding Penicillin-Binding Protein 7 of Escherichia Coli.” Journal of Bacteriology 177 (8): 2074–79.

Hong, Samuel, Albert Rovira, Peter Davies, Christina Ahlstrom, Petra Muellner, Aaron Rendahl, Karen Olsen, et al. 2016. “Serotypes and Antimicrobial Resistance in Salmonella Enterica Recovered from Clinical Samples from Cattle and Swine in Minnesota, 2006 to 2015.” PLOS ONE 11 (12): e0168016. https://doi.org/10.1371/journal.pone.0168016.

Jaffe, A, Y A Chabbert, and O Semonin. 1982. “Role of Porin Proteins OmpF and OmpC in the Permeation of Beta-Lactams.” Antimicrobial Agents and Chemotherapy 22 (6): 942–48.

Klemm, Elizabeth J., Sadia Shakoor, Andrew J. Page, Farah Naz Qamar, Kim Judge, Dania K. Saeed, Vanessa K. Wong, et al. 2018. “Emergence of an Extensively Drug-Resistant Salmonella Enterica Serovar Typhi Clone Harboring a Promiscuous Plasmid Encoding Resistance to Fluoroquinolones and Third-Generation Cephalosporins.” mBio 9 (1): e00105–18. https://doi.org/10.1128/mBio.00105-18.

Korsak, Dorota, Sylvia Liebscher, and Waldemar Vollmer. 2005. “Susceptibility to Antibiotics and β-Lactamase Induction in Murein Hydrolase Mutants of Escherichia Coli.” Antimicrobial Agents and Chemotherapy 49 (4): 1404–9. https://doi.org/10.1128/AAC.49.4.1404-1409.2005.

Krishnamoorthy, Ganesh, Elena B. Tikhonova, Girija Dhamdhere, and Helen I. Zgurskaya. 2013. “On the Role of TolC in Multidrug Efflux: The Function and Assembly of AcrAB-TolC Tolerate Significant Depletion of Intracellular TolC Protein.” Molecular Microbiology 87 (5): 982–97. https://doi.org/10.1111/mmi.12143.

Krishnamoorthy, Ganesh, David Wolloscheck, Jon W. Weeks, Cameron Croft, Valentin V. Rybenkov, and Helen I. Zgurskaya. 2016. “Breaking the Permeability Barrier of Escherichia Coli by Controlled Hyperporination of the Outer Membrane.” Antimicrobial Agents and Chemotherapy 60 (12): 7372–81. https://doi.org/10.1128/AAC.01882-16.

Laemmli, U. K. 1970. “Cleavage of Structural Proteins during the Assembly of the Head of Bacteriophage T4.” Nature 227 (5259): 680–85.

Matsuhashi, Michio, Yohtaroh Takagaki, Ichiro N. Maruyama, Shigeo Tamaki, Yukinobu Nishimura, Hideho Suzuki, Utako Ogino, and Yukinori Hirota. 1977. “Mutants of Escherichia Coli Lacking in Highly Penicillin-Sensitive D-Alanine Carboxypeptidase Activity.” Proceedings of the National Academy of Sciences of the United States of America 74 (7): 2976–79.

Matsuhashi, M, S Tamaki, S J Curtis, and J L Strominger. 1979. “Mutational Evidence for Identity of Penicillin-Binding Protein 5 in Escherichia Coli with the Major D-Alanine Carboxypeptidase IA Activity.” Journal of Bacteriology 137 (1): 644–47.

Meberg, Bernadette M., Avery L. Paulson, Richa Priyadarshini, and Kevin D. Young. 2004. “Endopeptidase Penicillin-Binding Proteins 4 and 7 Play Auxiliary Roles in Determining Uniform Morphology of Escherichia Coli.” Journal of Bacteriology 186 (24): 8326–36. https://doi.org/10.1128/JB.186.24.8326-8336.2004.

Miller, Jeffrey H. 1972. Experiments in Molecular Genetics. Cold Spring Harbor Laboratory.

Nikaido, H., and M. Vaara. 1985. “Molecular Basis of Bacterial Outer Membrane Permeability.” Microbiological Reviews 49 (1): 1–32.

Nishimura, Y, H Suzuki, Y Hirota, and J T Park. 1980. “A Mutant of Escherichia Coli Defective in Penicillin-Binding Protein 5 and Lacking D-Alanine Carboxypeptidase IA.” Journal of Bacteriology 143 (1): 531–34.

Ofek, I., S. Cohen, R. Rahmani, K. Kabha, D. Tamarkin, Y. Herzig, and E. Rubinstein. 1994. “Antibacterial Synergism of Polymyxin B Nonapeptide and Hydrophobic Antibiotics in Experimental Gram-Negative Infections in Mice.” Antimicrobial Agents and Chemotherapy 38 (2): 374–77. https://doi.org/10.1128/AAC.38.2.374.

Prestidge, Louise S., and Arthur B. Pardee. 1957. “INDUCTION OF BACTERIAL LYSIS BY PENICILLIN1.” Journal of Bacteriology 74 (1): 48–59.

Rhen, Mikael, C. David O’Connor, and Soila Sukupolvi. 1988. “The Outer Membrane Permeability Mutation of the Virulence-Associated Plasmid of Salmonella Typhimurium Is Located in a traT-like Gene.” FEMS Microbiology Letters 52 (1-2): 145–53. https://doi.org/10.1111/j.1574-6968.1988.tb02586.x.

Sabet, S. F., and C. A. Schnaitman. 1973. “Purification and Properties of the Colicin E3 Receptor of Escherichia Coli.” The Journal of Biological Chemistry 248 (5): 1797–1806.

Sauvage, Eric, Frédéric Kerff, Mohammed Terrak, Juan A. Ayala, and Paulette Charlier. 2008. “The Penicillin-Binding Proteins: Structure and Role in Peptidoglycan Biosynthesis.” FEMS Microbiology Reviews 32 (2): 234–58. https://doi.org/10.1111/j.1574-6976.2008.00105.x.

Schmieger, H. 1972. “Phage P22-Mutants with Increased or Decreased Transduction Abilities.” Molecular & General Genetics: MGG 119 (1): 75–88.

Singh, Santosh Kumar, Sadiya Parveen, L. SaiSree, and Manjula Reddy. 2015. “Regulated Proteolysis of a Cross-Link–specific Peptidoglycan Hydrolase Contributes to Bacterial Morphogenesis.” Proceedings of the National Academy of Sciences 112 (35): 10956–61. https://doi.org/10.1073/pnas.1507760112.

Singh, Santosh Kumar, L. SaiSree, Ravi N. Amrutha, and Manjula Reddy. 2012. “Three Redundant Murein Endopeptidases Catalyse an Essential Cleavage Step in Peptidoglycan Synthesis of Escherichia coliK12.” Molecular Microbiology 86 (5): 1036–51. https://doi.org/10.1111/mmi.12058.

Spratt, B G. 1980. “Deletion of the Penicillin-Binding Protein 5 Gene of Escherichia Coli.” Journal of Bacteriology 144 (3): 1190–92.

Sukupolvi, S, A Edelstein, M Rhen, S J Normark, and J D Pfeifer. 1997. “Development of a Murine Model of Chronic Salmonella Infection.” Infection and Immunity 65 (2): 838–42.

Sukupolvi, S., M. Vaara, I. M. Helander, P. Viljanen, and P. H. Mäkelä. 1984. “New Salmonella Typhimurium Mutants with Altered Outer Membrane Permeability.” Journal of Bacteriology 159 (2): 704–12.

Tadokoro, Akiko, Hidemi Hayashi, Toshihiko Kishimoto, Yasutaka Makino, Shingo Fujisaki, and Yukinobu Nishimura. 2004. “Interaction of the Escherichia Coli Lipoprotein NlpI with Periplasmic Prc (Tsp) Protease.” Journal of Biochemistry 135 (2): 185–91.

Taira, S., P. Riikonen, H. Saarilahti, S. Sukupolvi, and M. Rhen. 1991. “The mkaC Virulence Gene of the Salmonella Serovar Typhimurium 96 Kb Plasmid Encodes a Transcriptional Activator.” Molecular & General Genetics : MGG 228 (3): 381–84. https://doi.org/10.1007/BF00260630.

Vaara, M., and T. Vaara. 1983. “Sensitization of Gram-Negative Bacteria to Antibiotics and Complement by a Nontoxic Oligopeptide.” Nature 303 (5917): 526–28.

Weeks, Jon W., Teresa Celaya-Kolb, Sara Pecora, and Rajeev Misra. 2010. “AcrA Suppressor Alterations Reverse the Drug Hypersensitivity Phenotype of a TolC Mutant by Inducing TolC Aperture Opening.” Molecular Microbiology 75 (6): 1468–83. https://doi.org/10.1111/j.1365-2958.2010.07068.x.

Yu, D., H. M. Ellis, E. C. Lee, N. A. Jenkins, N. G. Copeland, and D. L. Court. 2000. “An Efficient Recombination System for Chromosome Engineering in Escherichia Coli.” Proceedings of the National Academy of Sciences of the United States of America 97 (11): 5978–83. https://doi.org/10.1073/pnas.100127597.

